# AlphaFold Model Quality Self-Assessment Improvement Via Deep Graph Learning

**DOI:** 10.1101/2024.07.18.604136

**Authors:** Jacob Verburgt, Zicong Zhang, Daisuke Kihara

## Abstract

In the past several years significant advances have been made in the field of deep-learning based computational modeling of proteins, with DeepMind’s AlphaFold2 being among the most prominent. Alongside the atom coordinates, these computationally modeled protein structures typically contain self-confidence metrics that can be used to gauge the relative modeling quality of individual residues, or the protein as a whole. Unfortunately, these scores are not always accurate, and may sometimes annotate poorly modeled regions of the protein as high confidence. Here, we introduce EQA-Fold to address this problem. EQA-Fold overhauls the LDDT prediction head of AlphaFold to provide more accurate self-confidence scores. We show that EQA-Fold is able to provide more accurate self-confidence scores than the standard AlphaFold architectures, as well as recent Model Quality Assessment protocols.

## Introduction

With the introduction of AlphaFold2 [1] in the CASP14 [2] protein structure modeling competition, the field of protein structure prediction rapidly matured, and a copious amount of computationally predicted protein structures became available through online databases such as the AlphaFold Protein Structure Database [3] and the Protein Structure Databank (PDB). Along with 3D coordinates of the atoms within each of these predicted structures, each residue is also annotated with a self-confidence score, which are used to gauge how trustworthy the modeling of that region of the structure is. Within the context of deep learning based methods, these confidence scores are often predictions of traditional quality metrics that have been used in the field of protein structure prediction for many years, with the underlying prediction network being trained in conjunction with the structure prediction itself. The primary confidence metric predicted within AlphaFold2 is the Local Distance Difference Test (LDDT) [4], which is predicted for each residue. These confidence scores are often overlooked or sidelined to the underlying structure prediction, but are often critical for downstream applications researchers in gauging the reliability of structures. Furthermore, in the event that multiple structure predictions exist, as they often do, these confidence scores are used to rank and select them, bolstering the significance of the self-confidence scores for downstream research.

In a similar vein, there are a number of Model Quality Assessment (MQA) methods that have been developed to assign quality scores to already predicted protein structures [5], [6], [7], [8]. Instead of relying on self-confidence metrics generated alongside the predicted models, these methods instead analyze the resultant protein structures to assign their own confidence scores. This allows for the analysis of protein structures generated from a variety of methods, which is often of use for the CASP Estimation of Model Accuracy category [9], in which participants attempt to rank and select well-modeled structures from a pool of candidate structures. The graph-like nature of the LDDT metric has led many of these methods to be graph-based [5], [7], [8].

In this study, we aim to increase the accuracy of AlphaFold2 like pLDDT self-confidence scores through incorporation of equivariant graph neural networks (EGNNs) [10] in place of the standard LDDT prediction head present within AlphaFold. EGNN architectures are able to leverage relative spatial information within the graph and are widely used for parsing and interpretation of molecular data [8], [11], [12], including MQA methods [13], and have been shown to outperform traditional graph methods in similar tasks [8]. Titled Equivariant Quality Assessment Folding (EQA-Fold), this work reimplements and fine-tunes the LDDT prediction head of a pre-trained AlphaFold model to provide more accurate self-confidence scores.

This process differs substantially from standard MQA methods by instead improving the underlying self-confidence scores that are generated alongside the predicted structure, as opposed to applying an external program for analysis. This provides us with the unique benefit of leveraging the same information that was used to generate the predicted protein structure. We believe that improvement in self-confidence score accuracy can be achieved through several key concepts:

1. EGNN architecture will allow better pLDDT assignment over the multi-layer perceptron network used within AlphaFold. In particular, the AlphaFold pLDDT prediction head does not leverage pairwise information, which we will leverage as edges, as well as edge features within our network.
2. Cleaning of fine-tuning data. Within the training of AlphaFold, and more broadly structure prediction, it is common to extract single protein chains out of multimeric structures. Although this can provide more data for training, it comes at severe detriment to the quality of the training data.
3. Incorporation of additional data. In addition to incorporating the pairwise information present within the AlphaFold framework, we also leverage features generated outside of AlphaFold. In particular, we utilize fluctuations that occur when the model is ran multiple times, and embedding data from protein language models.

## Materials and Methods

### Dataset

We derive our datasets from the precomputed data from the PISCES protein sequence culling server [14]. Primarily, we use data pulled from the December 14, 2020 release, culled at 90%, and have a resolution of at least 2.5 Å. These protein chains were fetched from the PDB, and renumbered to match their respective FASTA sequence, resulting in a total of 33,635 protein chains. For finetuning the LDDT prediction head, we further filtered our dataset to only contain protein chains where each chain exists as a standalone protein. We do this by fetching the equivalent biological assembly of the PDB entry of the chain, and filtering out models where multiple chains were present within the biological assembly, and had a minimum of 40 resolved residues. This resulted in a subset of 11,966 single-chain proteins.

The primary validation data was pulled from the PDB on March 1^st^, 2022. The entire PDB was filtered to find chains derived from X-ray structures with 2.5 Å or higher resolution and contained at least 20 residues in the sequence. These sequence chains were then clustered against the previous PISCES data via MMseqs2 [15] with a 90% similarity cutoff. These protein chains were fetched from the PDB, and renumbered to match their respective FASTA sequence, resulting in a total of 364 structures after feature generation.

We generate our testing dataset from the May 31^st^, 2023 release of the PISCES protein sequence culling server with the same constraints. To remove redundancy with the previously collected data, we first removed entries with identical PDB ids, and subsequently clustered the respective sequences against the previous PISCES and validation data via MMseqs2 [15] with a 90% similarity cutoff. These protein chains were fetched from the PDB, and renumbered to match their respective FASTA sequence, resulting in a total of 3,582 structures after feature generation. We again further filtered our testing dataset to only contain protein chains where each chain exists as a standalone protein by referencing the biological assembly, and had a minimum of 40 resolved residues, resulting in a total of 726 single-chain proteins

### Feature Generation

For all data, we generate AlphaFold “Single” (LX384) and “Pair” (LXLX128) embeddings using the AlphaFold 2.1 release, in which L is the length of the sequence. We precomputed MSAs for all targets using HHblits [16] and HHsearch [17] using the UniRef30 and PDB70 databases. Embeddings are output from the AlphaFold evoformer architecture, and contain information about individual residues, as well as information between residue pairs respectively. Furthermore, we generate Evolutionary Scale Modeling (ESM) embeddings using the esm2_t33_650M_UR50D model [18]. For each chain, we generate the sequence embedding (LX1280), the final attentions (LXLX33X20), as well as the layer-wise averages for the embeddings and attentions (LX33), (LXLX33) as described in the EquiPPIS protocol [8].

To leverage data from structural variability, we also run the standard AlphaFold architecture for 5 replications with a 50% dropout rate in all structure module weights. The resultant 5 models are then structurally superimposed, and a Cα Root Mean Square Fluctuation (RMSF) value is subsequently computed for each residue. These RMSF values are then subsequently used as residue level features in the LDDT prediction network. Our motivation for incorporating RMSF values stems from consensus methods for protein structure prediction and analysis as seen in previous CASP rounds, as well as previously reported benefits from enhanced sampling from AlphaFold [19]. Furthermore, previous studies have shown that pLDDT values correlate well with RMSF values calculated from molecular dynamics trajectories [20].

### Network Architecture

We primarily base our method off of our in-house implementation of AlphaFold, which is built on a composite of OpenFold [21] and newly generated code. Our method starts at the structure module stage of AlphaFold, with the Single and Pair representations being precomputed via the DeepMind AlphaFold implementation.

The newly developed LDDT prediction head converts the final single and pair representations, predicted Cα coordinates, and ESM layer-wise embeddings into a graph representation (**Figure 1**). Node features represent residues and are constructed by concatenating the final single representation (LX384) and the averaged ESM layers (LX33) and passing through a linear transition layer with ReLU activations, resulting in LX384 node features (**Figure 1A**). Edges are constructed for residues in which Cα positions were within 16Å. Edge features are constructed using the pair embeddings for any given residue. For any edge, the features from the pairwise embedding (LXLX128) for that residue pair are extracted and used as edge features (**Figure 1B**). The EGNN based prediction network itself consists of 4 equivariant graph convolutional layers with 384 node features at the input, 128 hidden node features between layers and 50 output node features.

**Figure 1:**
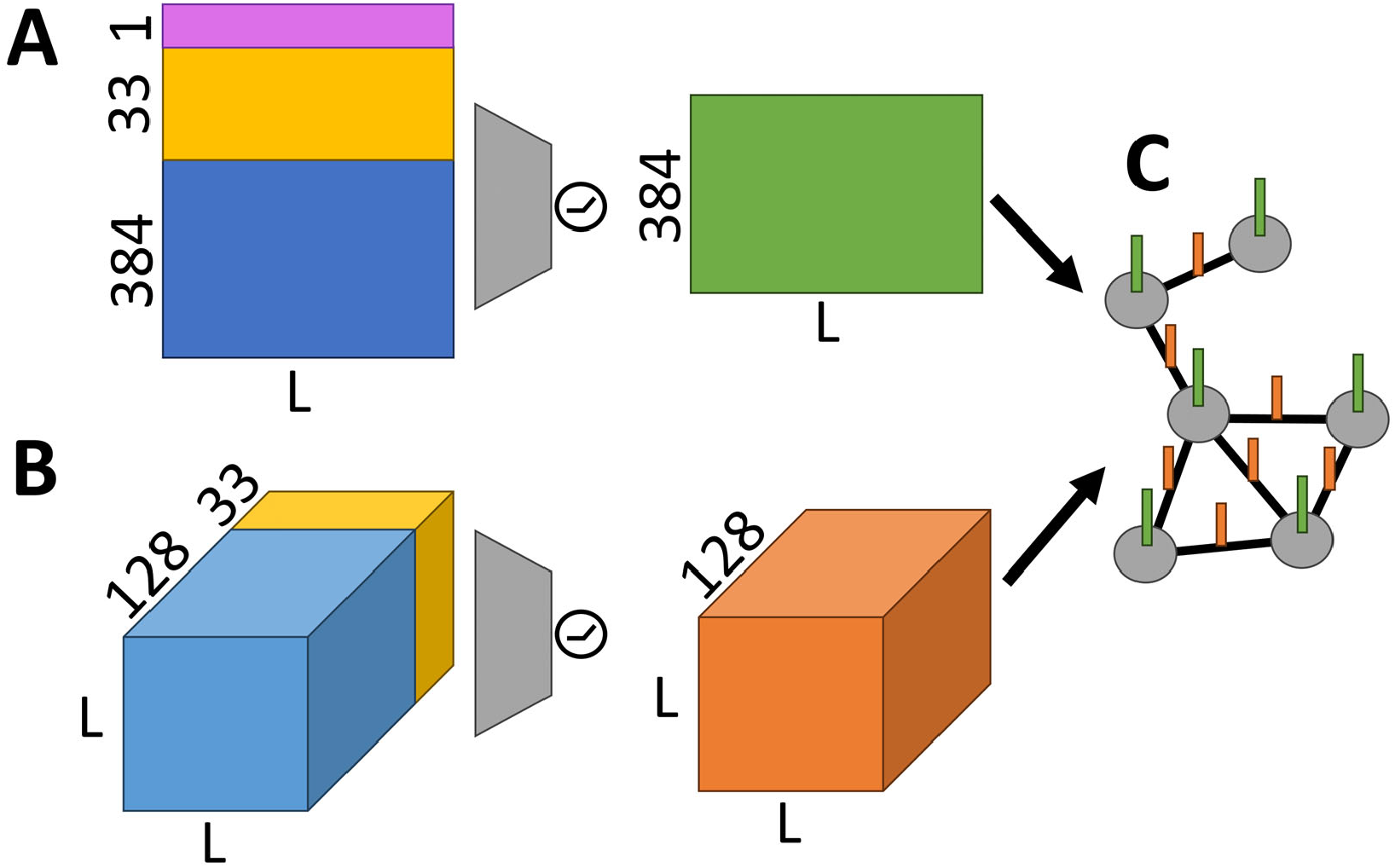
Construction of the Protein Graph Representation for the LDDT Prediction Head. A) The final single representation (blue), layerwise ESM embeddings (yellow), and RMSF values (magenta) are concatenated and passed through a single linear transition layer with ReLU activation and are subsequently used as node features. B) AlphaFold Pairwise embeddings (blue) and ESM Attentions (yellow) are passed through a single transition layer with ReLU activation, contain information for all residue pairs, which are extracted as edge features. C) The graph representation of the protein is built using Cα distances and has the constructed node and edge features loaded on.

### Training and Implementation

The base network is built and trained with PyTorch 1.12.0, with graph operations being done in PyTorch-Geometric 2.3.0 and the EGNN implementation from the nascent paper from Satorras *et al*. [10].

The full network, with the standard AlphaFold pLDDT prediction head, was initially trained with a learning rate of 1 e-3, and a batch size of 4 using the 30K training dataset, with a network architecture identical to what is described in the AlphaFold2 protocol. After initial training, all structure module parameters were frozen, and the standard LDDT prediction head was replaced by the EGNN model.

Furthermore, we modify the loss function by weighting the LDDT residuals that are generated within the pLDDT loss function. Since the highly accurate models largely skew the LDDT distribution towards high values, we account for this by first calculating relative weights for each of the 50 LDDT bins. The weight for any LDDT bin is equal to 1 – the relative fraction of LDDT values that fall within that bin in the training data. In practice, this down-weights and corrects for the overrepresented LDDT values in the loss calculation.

### Analysis

The Local Distance Difference Test (LDDT) score provides a superimposition-free metric to gauge the relative modeling quality of a predicted protein structure to that of an experimentally resolved ground-truth structure [4]. Given a residue in a predicted structure and an experimentally determined ground-truth structure, the LDDT score looks at all Cα atoms within a 15 Å inclusion proximity within the Cα atom of the residue of interest in the true structure. For each of these Cα atom pairs, the relative distance between the ground-truth and the predicted structures are compared. Specifically, the score checks if the differences in the distances are within 0.5 Å, 1 Å, 2 Å, and 4 Å of each other. A Cα atom pair can meet all thresholds, (resulting in a score of 1), or none of the thresholds (resulting in a score of 0). This is repeated and averaged for all Cα atom pairs within the inclusion proximity around the protein of interest to generate an LDDT score for that residue and is subsequently repeated for all residues within the protein. Alternatively, all heavy atoms of the residue of interest can be considered, resulting in a different, and more stringent score. We term this the “all-atom LDDT” (LDDT-AA) score. These scores are often scaled between 0 and 100 when annotating predicted structures.

Ground truth LDDT values were calculated using the OpenStructure executable [4], [22] version 2.3.1. For the standard Cα only LDDT values, the “-c” flag was used to only consider Cα atoms. To compute all-atom LDDT values (LDDT-AA), no flag was used to consider all heavy atoms. We primarily gauge performance of LDDT assignment by computing the LDDT error for each target in the test dataset. To compute the error, the LDDT error for any given residue i is computed by taking the absolute difference between the predicted LDDT value, and the true LDDT value (Equation 1). These values are then averaged for all resolved residues in in the protein model to compute the LDDT error (Equation 2). Results are visualized in PyMOL version 2.4 [23].

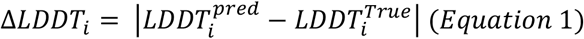

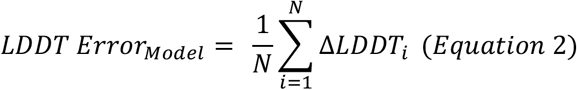

### Alternate Methods

#### AlphaFold DataBase

For every target within the test dataset, we collect the deposited AlphaFold DataBase (AFDB) [3] model via the web API by cross referencing UniProt IDs [24] between the PDB entry and AFDB, for entries in which a Uniprot ID was available, and an AFDB entry was present for that ID. We renumber the AFDB model to match the residue numbering used in the testing dataset. In total, 530 of the 726 test set structures had AFDB entries successfully retrieved and formatted.

## Results

### Overall Results

To analyze the performance of our models, we calculate the model-level LDDT error for each target in the test set for both EQA-Fold and the standard AlphaFold architecture (**Figure 2A**). From the 726 targets in the Test set, EQA-Fold was able to outperform the standard AlphaFold architecture for 111 (15.29%) of the targets, with 582 (80.17%) being “tied” within a margin of 0.005. The standard AlphaFold Architecture only outperformed EQA-Fold in 33 (4.55%) targets.

**Figure 2:**
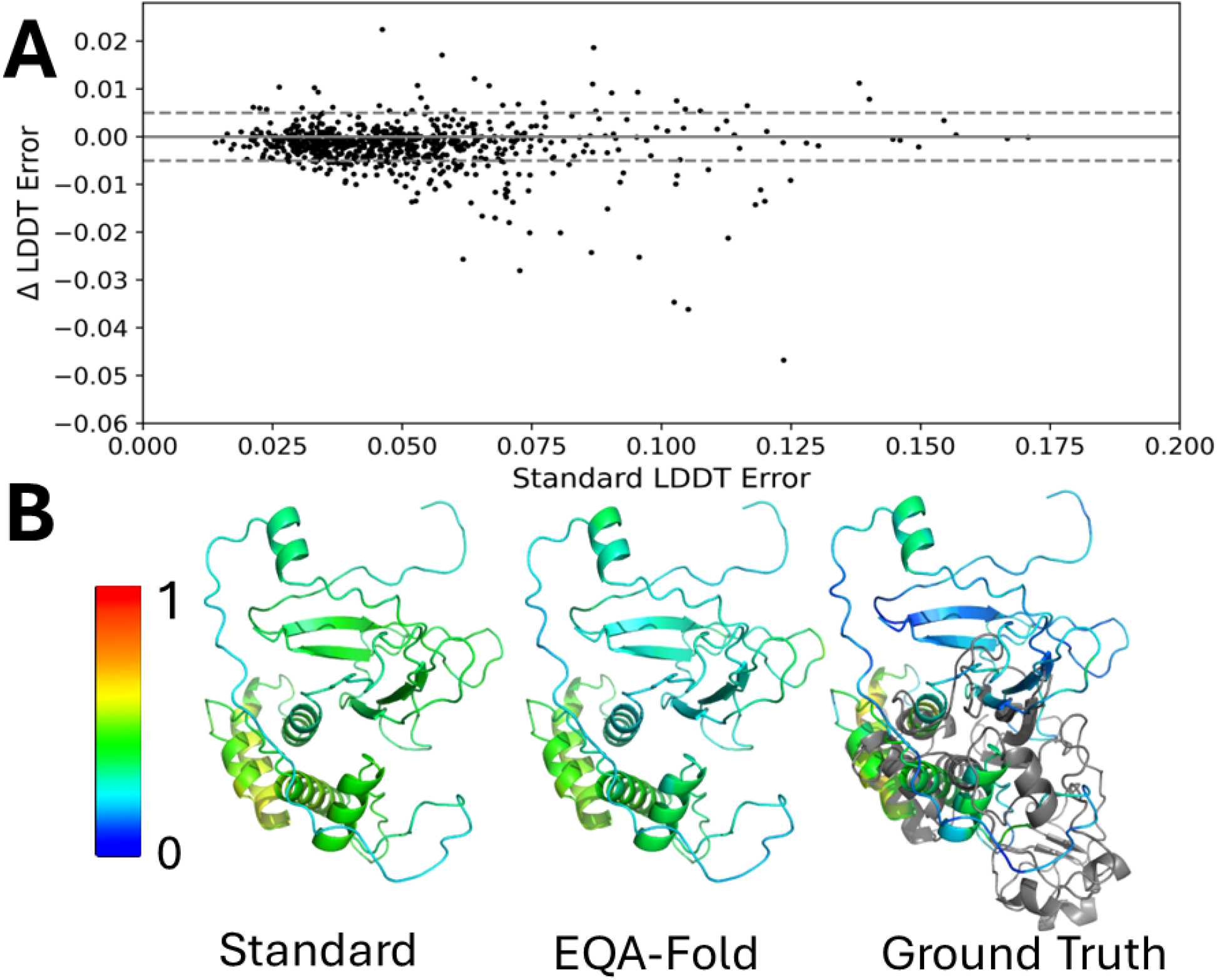
Model Quality Assignment Improvement by EQA-Fold. A) Deviations in average LDDT error the standard AlphaFold architecture models and EQA-Fold on all targets present within the test dataset. The error for any given model, the error is calculated by averaging the absolute LDDT error for any given residue for all residues in the model. B) Visualization of LDDT improvement for test set target c. The standard architecture model (top, left) and the EQA-Fold model (top, right), and true LDDT values (bottom) are colored from Blue (pLDDT of 0, to red (pLDDT of 100). The experimentally determined ground truth structure is shown in gray.

However, it is important to remember that both EQA-Fold and AlphaFold generally perform well in determining self-confidence scores. Since we are particularly interested in the models with a large amount of LDDT error, we look at the subset of the data where either method had an average LDDT error of above 0.05. Of the 301 models that meet this criteria, EQA-Fold outperforms the standard AlphaFold architecture for 75 (24.92%) of the targets, with 201 (66.78%) targets being within the “tied” margin. The standard AlphaFold architecture only had lower LDDT Error than EQA-Fold for 25 (8.31%) targets within this subset.

We visualize the deviations in pLDDT assignment between EQA-Fold and the standard AlphaFold architecture for the Rubella Protease (PDB: 7FAV) in **Figure 2B**. The AlphaFold structure module modeled the C-terminal region poorly (LDDT values between 0.095-0.206). For these residues, the standard AlphaFold architecture annotated these residues with pLDDT values in the range of 0.334-0.402. EQA-Fold improves these values by labeling them as less-confident, and closer to the true LDDT range (pLDDT values 0.252-0.284.

### Benchmarking against other methods

We compared the accuracy of the self confidence scores produced by EQAFold models to those of AFDB entries to gauge the relative accuracy of LDDT modeling.

#### AlphaFold DB

For each target in the test set, we fetched the equivalent model from AFDB, provided an equivalent entry exists, via the UniProt ID (see methods). Out of the 726 targets in the test set, we were able to identify and clean AFDB entries for 530 of them. For each residue in each target, we computed the error in LDDT as the absolute difference between true and predicted LDDT values (**Figure 3A**). Out of the 530 targets, EQA-Fold had less overall LDDT error in 159 (30.00%) of cases, whereas AFDB was able to maintain lower error in 179 (33.77%) of models. 192 (36.23%) of models were within the “tied” margin of error. However, when focusing on the 256 targets where either method had an average LDDT error of above 0.05, EQA-Fold outperformed the Deposited AFDB model in 127 (49.61%) of targets, with 56 (21.88%) of models being tied. Only 73 (28.52%) of AFDB models with substantial modeling error had lower LDDT error than EQA-Fold.

**Figure 3:**
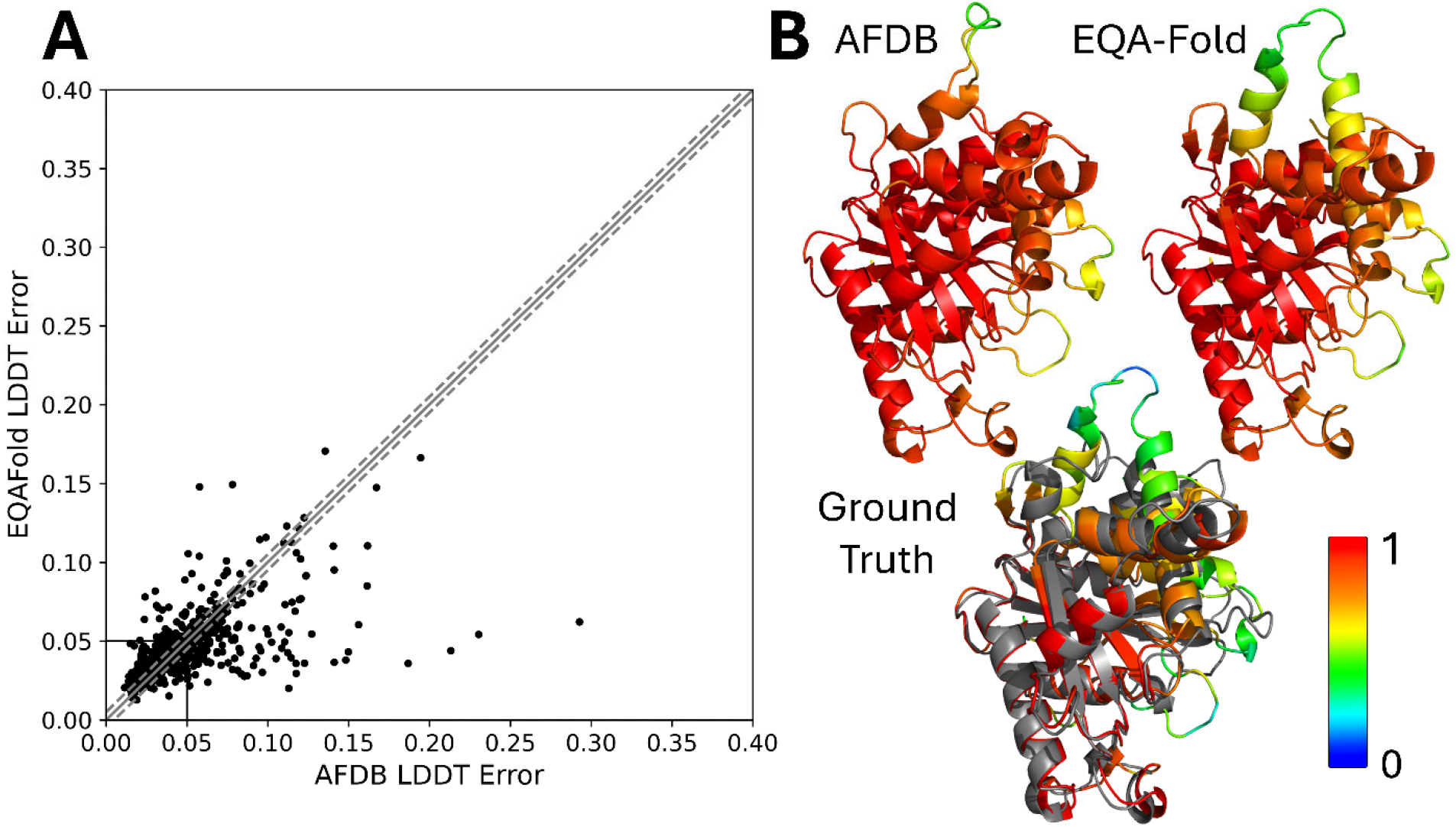
Relative LDDT Prediction Performance Compared to AFDB models. **A)** Deviations in average LDDT error between deposited AFDB models and EQA-Fold on all targets present within the test dataset. The error for any given model, the error is calculated by averaging the absolute LDDT error for any given residue for all residues in the model. Each black point represents a target in the test set. The black box indicates the boundary in which both methods had substantial (>0.05) LDDT error. The solid gray line represents the y=x line, and dashed gray lines represent the boundary in which LDDT errors are considered equivalent. **B)** Visualization of LDDT improvement for test set target 6VKJ. The AFDB model (top, left) and the EQA-Fold model (top, right), and true LDDT values (bottom) are colored from Blue (pLDDT of 0, to red (pLDDT of 1). The experimentally determined ground truth structure is shown in gray.

We visualize the quality of AFDB LDDT assignments against those of EQA-Fold in **Figure 3B**. LDDT assignment for Human Guanylate Binding Protein 2 (PDB: 6VKJ). The structure prediction for this target was generally accurate for both AFDB and EQA-Fold. The major deviation from the ground-truth structure for both AFDB and EQA-Fold is the improperly modeled loop region. The helix leading up to this loop region has true LDDT values ranging between 0.367 and 0.716. AFDB annotates the helix leading up to this loop with predicted LDDT values ranging between 0.818-0.913. However, the poor modeling in this region is better realized in EQA-Fold, which properly labeled this region as low quality with predicted LDDT values ranging between 0.490-0.693.

#### EQA-Fold-AA

We train a variant if EQA-Fold to instead predict the all-atom LDDT (LDDT-AA) (**Figure 4A**). This metric considers all heavy atoms (not just Cα atoms) within the inclusion radius for any given residue. In practice, this makes the LDDT-AA metric a more stringent version of the LDDT metric (**Figure 4B**). Although this is a different metric than what AlphaFold predicts, it can be useful for identifying poorly modeled sidechain atoms and contacts. In particular, it can be useful for discerning small modeling errors in regions where the LDDT metric is nearly perfect (**Figure 4B, inset**). This prompted us to retrain EQA-Fold to predict the more stringent LDDT-AA metric. We facilitate this training by modifying the LDDT-Loss function to consider all-atoms between the true and predicted structures when calculating the true LDDT values.

**Figure 4:**
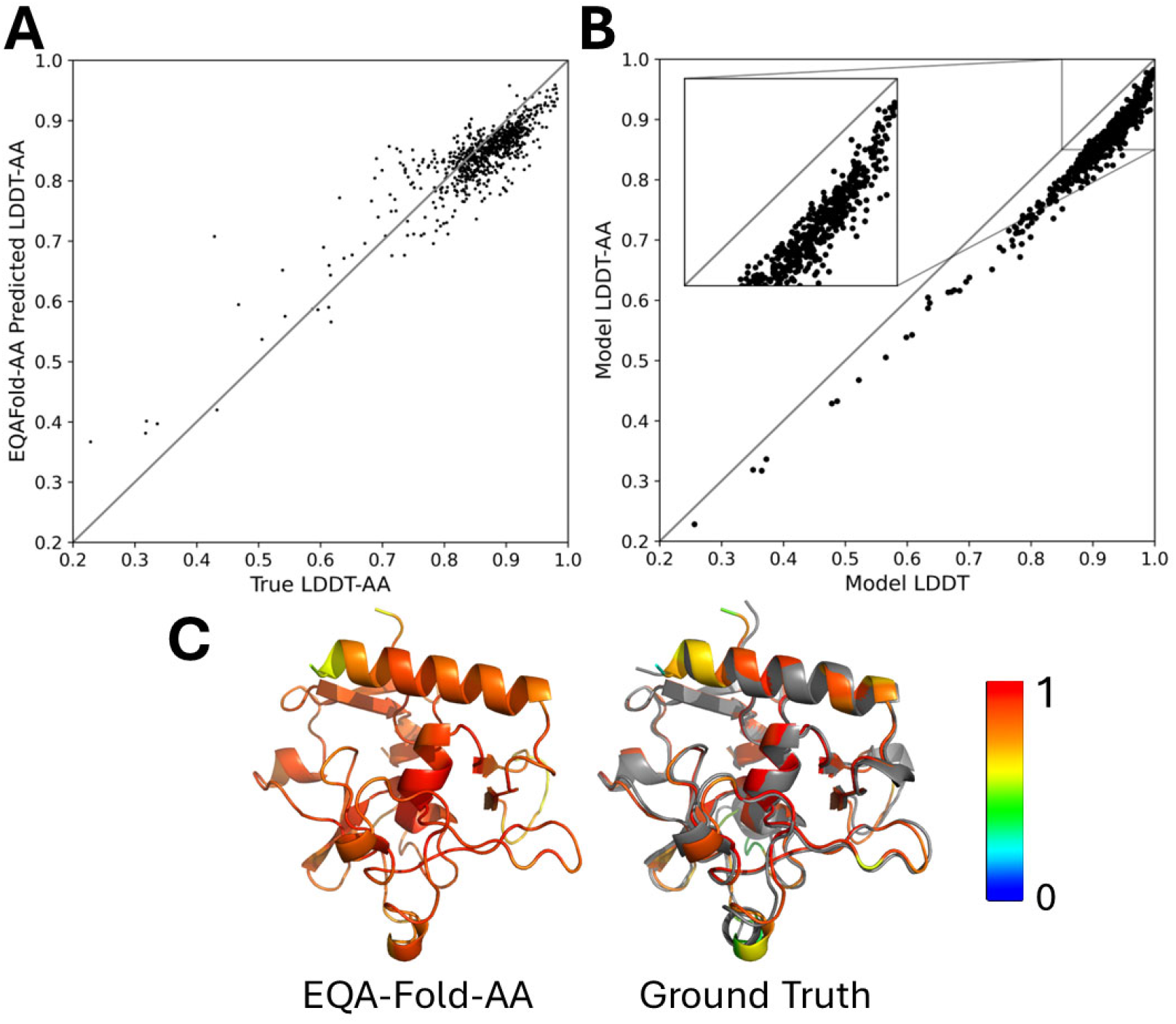
EQA-Fold-AA. **A)** Deviations in average LDDT-AA predictions by EQA-Fold-AA. The error for any given model, the error is calculated by averaging the absolute LDDT error for any given residue for all residues in the model. Each black point represents a target in the test set. **B)** Correlation between LDDT and LDDT-AA. For each target within the test set, the model LDDT and LDDT-AA values are plotted against each other. **D)** Visualization of LDDT-AA assignment for test set target 7D86. The EQA-Fold-AA model (left), and true LDDT-AA values (right) are colored from Blue (pLDDT-AA of 0, to red (pLDDT-AA of 1). The experimentally determined ground truth structure is shown in gray.

We visualize the results of LDDT-AA assignment for Zebrafish PHF14-PZP (PDB: 7D86) in **Figure 4D**. EQA-Fold-AA was able to accurately annotate the predicted structure with values close to the LDDT-AA scores. The helix at the core of the protein, which is modeled almost perfectly, has LDDT-AA values in the range of 0.907-0.968. This is accurately represented in the predicted LDDT-AA score as produced by EQA-Fold-AA, with values ranging between 0.887-0.945.

### Feature Analysis

Additional features were added to the model with the intent of providing additional information for the model. We included ESM embeddings as they have been successfully utilized in previous EGNN protocols [8], [13]. Furthermore, we intended on leveraging consensus data, as it previously been of use in many protein structure prediction related tasks, and RMSF values derived from molecualr dynamics correlations correlate well with LDDT values. These RMSF values derived from replicate dropout runs correlate well with the LDDT and pLDDT values (**Figure 5A**), making them rational to include within the pLDDT prediction network.

**Figure 5:**
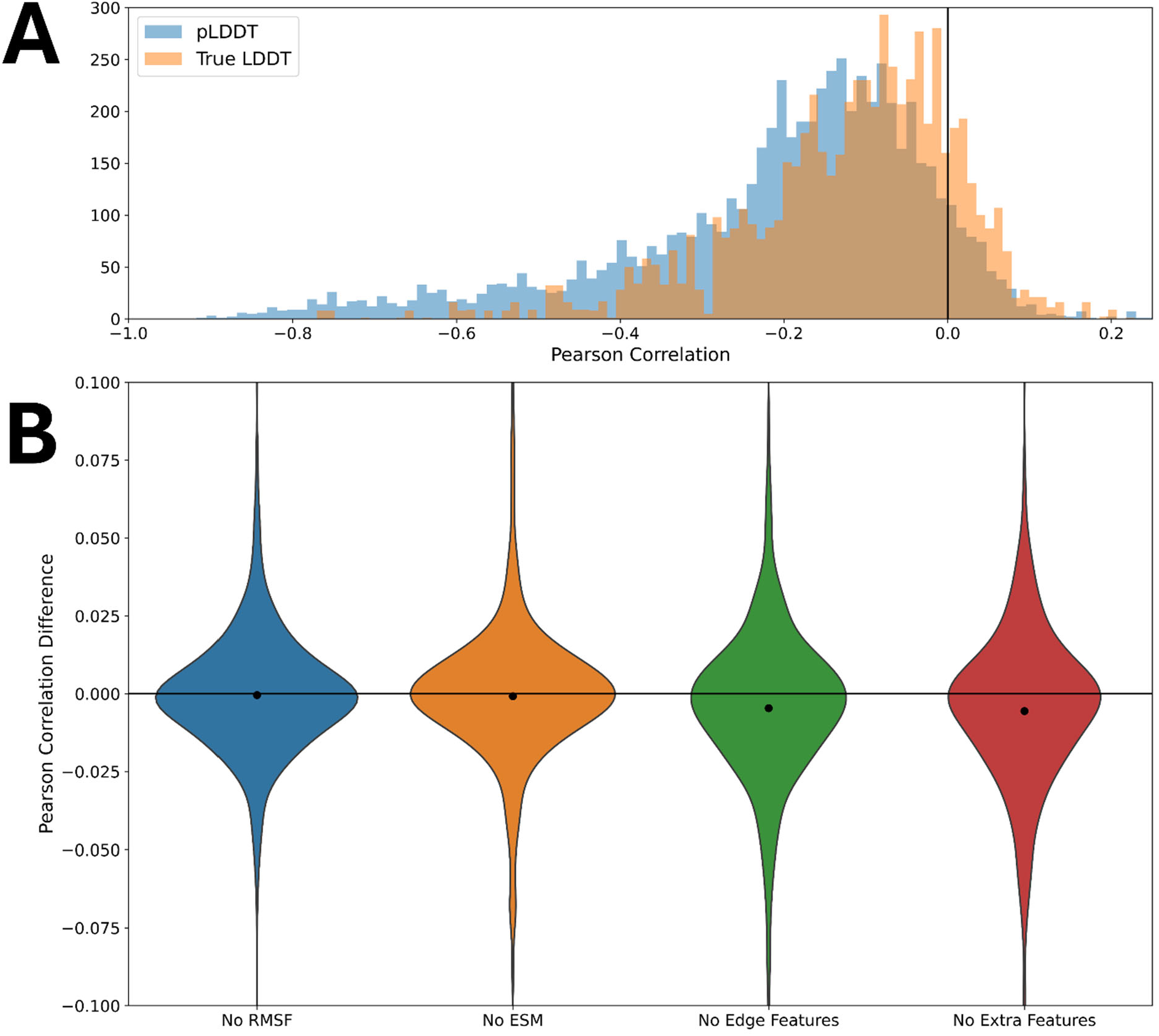
Effect of Sequential Removal of Features from EQA-Fold. **A)** Distribution of model Level correlations between RMSF and the true or predicted LDDT value. For each target within the test set, the RMSF value between all 5 dropout replicates was calculated for each residue, and compared to their respective LDDT or pLDDT values. Using all residues within the target, the Pearson correlation was calculated. The black line indicates the point of no correlation. **B)** The distribution of the Deviations of the Pearsons Correlation Coefficient from the full model for several model variants with different sets of features removed are shown. The black dot represents the average correlation for all models within the test set for that specific model.

To evaluate the relative contribution of node and edge features, we performed feature analysis by retraining several variant models with certain features, or sets of features omitted (**Figure 5B**). These variants include a model without any RMSF features (No RMSF), no ESM features (No ESM), no Edge Features (No Edge), and no additional features (No Extrafeat). For each model variant, we compute the Pearsons correlation coefficient for each target in our testing set by comparing the true LDDT values to the pLDDT values. **Figure 5** shows the relative decreases in model performance. The complete model had a correlation of 0.693. The removal of RMSF features (No RMSF) and ESM features (No ESM) resulted in almost no performance decreases of 0.692 and 0.692 respectively. The removal of all edge features (No Edge), and all additional features (No Extrafeat) had marginally lower correlations of 0.689 and 0.686 respectively. Although there are marginal increases in correlation when edge features are considered, it appears that the standard AlphaFold embeddings are sufficient for training the model.

## Discussion

Self-confidence metrics in protein structure prediction, such as the pLDDT score, are often overlooked components of protein structure prediction, but serve as critical metrics for the ranking and analysis of computationally predicted protein structures. This work introduced EQA-Fold and EQA-Fold-AA, which serve as AlphaFold-like architectures with improved self-confidence metric prediction architectures. The EGNN framework, explicitly embedded within the AlphaFold structure prediction architecture, is able to provide more accurate pLDDT annotations, thus increasing the reliability of predicted protein model self-confidence scores. Furthermore, since this method and protocol is present within, and trained alongside the protein structure prediction architecture, we believe that it is broadly applicable to a wide range of deep learning-based protein structure prediction methods. Future works on self-confidence improvement for protein structures may expand beyond tertiary structure prediction as explored here, and may cover cases such as multimeric complexes, multi-state models, and small molecule interactions.

## Acknowledgements

This work was partly supported by the National Institutes of Health (R01GM133840) and by the National Science Foundation (DBI2003635, DBI2146026, IIS2211598, DMS2151678, CMMI1825941, and MCB1925643).

## References

[1] J. Jumper et al., “Highly accurate protein structure prediction with AlphaFold,” Nature, vol. 596, no. 7873, pp. 583–589, Aug. 2021, doi: 10.1038/s41586-021-03819-2.

[2] M. F. Lensink et al., “Prediction of protein assemblies, the next frontier: The CASP14-CAPRI experiment,” Proteins: Structure, Function, and Bioinformatics, vol. 89, no. 12, pp. 1800–1823, 2021, doi: 10.1002/prot.26222.

[3] M. Varadi et al., “AlphaFold Protein Structure Database: massively expanding the structural coverage of protein-sequence space with high-accuracy models,” Nucleic Acids Res, vol. 50, no. D1, pp. D439–D444, Jan. 2022, doi: 10.1093/nar/gkab1061.

[4] V. Mariani, M. Biasini, A. Barbato, and T. Schwede, “lDDT: a local superposition-free score for comparing protein structures and models using distance difference tests,” Bioinformatics, vol. 29, no. 21, pp. 2722–2728, Nov. 2013, doi: 10.1093/bioinformatics/btt473.

[5] F. Baldassarre, D. Menéndez Hurtado, A. Elofsson, and H. Azizpour, “GraphQA: protein model quality assessment using graph convolutional networks,” Bioinformatics, vol. 37, no. 3, pp. 360–366, Feb. 2021, doi: 10.1093/bioinformatics/btaa714.

[6] K. Olechnovič and Č. Venclovas, “VoroMQA: Assessment of protein structure quality using interatomic contact areas,” Proteins: Structure, Function, and Bioinformatics, vol. 85, no. 6, pp. 1131–1145, 2017, doi: 10.1002/prot.25278.

[7] S. Roy and A. Ben-Hur, “Protein quality assessment with a loss function designed for high-quality decoys,” Front. Bioinform., vol. 3, Oct. 2023, doi: 10.3389/fbinf.2023.1198218.

[8] R. Roche, B. Moussad, M. H. Shuvo, and D. Bhattacharya, “E(3) equivariant graph neural networks for robust and accurate protein-protein interaction site prediction,” PLOS Computational Biology, vol. 19, no. 8, p. e1011435, Aug. 2023, doi: 10.1371/journal.pcbi.1011435.

[9] S. Kwon, J. Won, A. Kryshtafovych, and C. Seok, “Assessment of Protein Model Structure Accuracy Estimation in CASP14: Old and New Challenges,” Proteins, vol. 89, no. 12, pp. 1940–1948, Dec. 2021, doi: 10.1002/prot.26192.

[10] V. G. Satorras, E. Hoogeboom, and M. Welling, “E(n) Equivariant Graph Neural Networks.” arXiv, Feb. 16, 2022. doi: 10.48550/arXiv.2102.09844.

[11] M. R. Masters, A. H. Mahmoud, Y. Wei, and M. A. Lill, “Deep Learning Model for Efficient Protein– Ligand Docking with Implicit Side-Chain Flexibility,” J. Chem. Inf. Model., vol. 63, no. 6, pp. 1695–1707, Mar. 2023, doi: 10.1021/acs.jcim.2c01436.

[12] T. Wu, Z. Guo, and J. Cheng, “Atomic protein structure refinement using all-atom graph representations and SE(3)-equivariant graph transformer,” Bioinformatics, vol. 39, no. 5, p. btad298, May 2023, doi: 10.1093/bioinformatics/btad298.

[13] C. Chen, X. Chen, A. Morehead, T. Wu, and J. Cheng, “3D-equivariant graph neural networks for protein model quality assessment,” Bioinformatics, vol. 39, no. 1, p. btad030, Jan. 2023, doi: 10.1093/bioinformatics/btad030.

[14] G. Wang and R. L. Dunbrack Jr, “PISCES: a protein sequence culling server,” Bioinformatics, vol. 19, no. 12, pp. 1589–1591, Aug. 2003, doi: 10.1093/bioinformatics/btg224.

[15] M. Steinegger and J. Söding, “MMseqs2 enables sensitive protein sequence searching for the analysis of massive data sets,” Nat Biotechnol, vol. 35, no. 11, Art. no. 11, Nov. 2017, doi: 10.1038/nbt.3988.

[16] M. Remmert, A. Biegert, A. Hauser, and J. Söding, “HHblits: lightning-fast iterative protein sequence searching by HMM-HMM alignment,” Nat Methods, vol. 9, no. 2, pp. 173–175, Feb. 2012, doi: 10.1038/nmeth.1818.

[17] J. Söding, “Protein homology detection by HMM–HMM comparison,” Bioinformatics, vol. 21, no. 7, pp. 951–960, Apr. 2005, doi: 10.1093/bioinformatics/bti125.

[18] Z. Lin et al., “Evolutionary-scale prediction of atomic-level protein structure with a language model,” Science, vol. 379, no. 6637, pp. 1123–1130, Mar. 2023, doi: 10.1126/science.ade2574.

[19] G. Brysbaert et al., “MassiveFold: unveiling AlphaFold’s hidden potential with optimized and parallelized massive sampling.” Apr. 30, 2024. doi: 10.21203/rs.3.rs-4319486/v1.

[20] H.-B. Guo et al., “AlphaFold2 models indicate that protein sequence determines both structure and dynamics,” Sci Rep, vol. 12, no. 1, p. 10696, Jun. 2022, doi: 10.1038/s41598-022-14382-9.

[21] G. Ahdritz et al., “OpenFold: Retraining AlphaFold2 yields new insights into its learning mechanisms and capacity for generalization.” bioRxiv, p. 2022.11.20.517210, Aug. 12, 2023. doi: 10.1101/2022.11.20.517210.

[22] M. Biasini et al., “OpenStructure: an integrated software framework for computational structural biology,” Acta Cryst D, vol. 69, no. 5, pp. 701–709, May 2013, doi: 10.1107/S0907444913007051.

[23] Schrödinger, LLC, “The PyMOL Molecular Graphics System, Version 1.8,” Nov. 2015.

[24] The UniProt Consortium, “UniProt: the Universal Protein Knowledgebase in 2023,” Nucleic Acids Research, vol. 51, no. D1, pp. D523–D531, Jan. 2023, doi: 10.1093/nar/gkac1052.

